# Endosymbiont strategic shifts inhibit cooperation during coral bleaching recovery

**DOI:** 10.1101/2022.06.28.497964

**Authors:** Luella Allen-Waller, Katie L. Barott

**Affiliations:** Department of Biology, University of Pennsylvania, Philadelphia, PA 19104 USA

## Abstract

The future of coral reefs in a warming world depends on corals’ ability to resist or recover from losing their photosynthetic algal endosymbionts (coral bleaching) during marine heatwaves. Heat-tolerant algal species can confer bleaching resistance by remaining in symbiosis during heat stress but tend to provide less photosynthate to the host than heat-sensitive species. Understanding this potential nutritional tradeoff is crucial for predicting coral success under climate change, but the energetic dynamics of corals hosting different algal species during bleaching recovery are poorly understood. To test how algal energetics affects coral recovery, we heat-stressed corals (*Montipora capitata*) hosting either heat-sensitive *Cladocopium* sp. or heat-tolerant *Durusdinium glynni* algae for two weeks, followed by a one-month recovery period. We found that while thermotolerant *D. glynni* regained density and photochemical efficiency faster after bleaching than *Cladocopium*, this algal recovery did not correspond with host physiological recovery, and *D. glynni* populations still contributed less photosynthate to the host relative to *Cladocopium*. Further, high-density algal populations of both species translocated a smaller proportion of their photosynthate than low-density populations, and corals receiving less photosynthate suffered reduced calcification rates and lower intracellular pH. This is the first evidence of a direct negative relationship between symbiont population size and ‘selfishness,’ and the first to establish a connection between Symbiodiniaceae carbon translocation and coral cellular homeostasis. Together, these results suggest that algal energy reallocation towards regrowth after bleaching can harm coral physiology, and that reestablishing a beneficial endosymbiosis can pose a secondary challenge for holobionts surviving stress.

## INTRODUCTION

Microbial endosymbiosis is a ubiquitous way of life that supports many key ecosystems [1, 2]. However, the benefits of many symbioses are context-dependent, with both environmental conditions and partner identity determining whether a particular association improves or impairs host fitness [3–5]. For example, while some endosymbionts have a neutral or negative effect on host fitness during stress, other symbiont species can confer stress tolerance on their hosts (e.g. [6–8]). Yet these hardy symbionts can also cost host fitness in the absence of that specific stress (e.g., bacteria that improve aphid heat tolerance but also reduce host fecundity relative to other bacterial strains [9], or fungal endophytes that enhance drought tolerance at the expense of host growth [10]). These patterns suggest that symbiotic stress tolerance (defined here as symbiont and host survival) may come at the expense of mutualistic function. This can produce a “hardiness-benefit” tradeoff, where hosts have either a stress-tolerant or a cooperative symbiont, the net benefits of which depend on environmental context. As global climate change outpaces genetic evolution for many species (e.g. [11, 12]), it is crucial to better understand how environmental disturbance will impact the persistence and function of mutualisms.

Reef-building corals rely on endosymbiotic dinoflagellate algae (Symbiodiniaceae) to survive, and the varying environmental sensitivity of this association may produce such a hardiness-benefit tradeoff. Under normal conditions, coral animals use energy gained from photosynthate released by their algal symbionts [13] to secrete the massive calcium carbonate skeletons that sustain reefs’ iconic biodiversity, including a staggering 25% of all marine species [14]. However, at just 1-2°C above mean summer temperatures, the coral-algal mutualism can break down in a process known as coral bleaching [15–17]. Some corals host Symbiodiniaceae species that remain *in hospite* despite high temperatures, possibly because these algae can maintain higher photochemical efficiency under heat stress than more thermosensitive symbionts ([18–21]; but see [22]). Corals with these thermotolerant algae are more likely to survive heat stress events (e.g. [23, 24]; but see [25]); however, these thermotolerant algae often provide less photosynthate to their hosts under ambient conditions, limiting host growth (e.g. [26–31]; but see [32, 33]). Coral symbionts may therefore exhibit a functional tradeoff between thermotolerance and nutritional benefit. It is critical to understand the consequences of this tradeoff for coral persistence as severe marine heatwaves increase in frequency and severity [34].

The energetic costs of recurrent bleaching may shift both the costs and benefits of symbiosis, potentially granting a crucial survival advantage to corals hosting thermotolerant symbionts despite their tendency to provide less fixed carbon [20, 30]. Reef futures depend on their corals’ ability to withstand increasingly frequent bleaching events, as bleached corals can starve within weeks if they do not regain their symbionts (e.g. [17, 23]), leaving reefs vulnerable to erosion [35, 36] and consequent biodiversity loss [37]. But while bleaching recovery requires the algal population to regrow [38, 39], host costs of symbiosis maintenance increase with both temperature and symbiont population density [40, 41] thus destabilizing symbiont cooperation [42–44]. Moreover, though colonization studies in coral juveniles suggest that thermotolerant symbionts grow faster than thermosensitive symbionts *in hospite* [45], we do not know whether this rapid growth supports symbionts’ ability to sustain host nutrition. Therefore, there is an urgent need to better understand Symbiodiniaceae photosynthate provisioning and use during bleaching recovery.

In order to examine how population recovery of different coral symbiont species affects host physiology after bleaching, we used a ‘living library’ of bleaching-susceptible and bleaching-resistant colonies of the reef-building coral *Montipora capitata* in Kāne’ohe Bay, Hawai’i [23]. These colonies have exhibited consistent bleaching phenotypes across multiple heatwaves [23, 46], and bleaching resistance has been attributed to the proportion of the thermotolerant symbiont *Durusdinium glynni* present: bleaching-resistant *M. capitata* tend to host primarily *D. glynni* (D-colonies), while bleaching-susceptible individuals host exclusively *Cladocopium* sp. (C-colonies) [47, 48]. Because these colonies were tagged in location- and depth-matched pairs that differ only in symbiont species and resulting bleaching phenotype, they are an ideal system to test for a symbiont hardiness-benefit tradeoff. Existing evidence already suggests C-colonies have a temperature-dependent advantage over D-colonies. Specifically, isotopic signatures indicate that these C-colonies assimilate more autotrophic carbon than D-colonies at ambient temperatures [31], suggesting that *Cladocopium* are more generous symbionts and matching trends for these symbiont genera in other regions. Meanwhile, a recent heatwave led to acidification of the intracellular pH (pH_i_) of C-colonies but not D-colonies, suggesting greater physiological stress in C-colonies during heat stress [46]. These changes in cellular homeostasis may be due to reductions in coral carbon assimilation in C-colonies during heat stress, which could be driven by disruptions in carbon translocation from the symbiont [42–44], or the loss of symbionts from the colony [46, 47]. However, what benefits corals hosting different symbiont species receive during and after heat stress remains unknown. Here we tested for the first time 1) whether heat-induced parasitism is generalizable across different dinoflagellate communities in the same environment, 2) how heat stress may alter coral carbon assimilation from different dinoflagellate communities in the same host species, and 3) whether variability in carbon assimilation drives differences in coral cellular homeostasis. This study is a novel and important step in understanding energy dynamics of different coral holobionts during bleaching recovery, and advances our understanding of how symbiotic partners allocate benefits in nutritional symbioses under stress.

## MATERIALS AND METHODS

### Field collections

Between June 7-10 2019, six fragments per genet (N = 60 fragments total) were sampled from previously identified bleaching-susceptible and bleaching-resistant depth-matched neighboring pairs of *M. capitata* colonies on Patch Reef 13, Kāne’ohe Bay, HI [23]. Corals were then transported to the Hawai’i Institute of Marine Biology in coolers containing ambient seawater. Immediately on arrival at HIMB, corals were placed in outdoor flow-through seawater tables. Water tables were semi-shaded to allow light levels similar to those on the reef (diel maximum of ~450 μmol m^-2^ sec^-1^ at 12:00). To keep corals upright, each was glued to a plastic base using non-toxic ethyl cyanoacrylate (CorAffix, Two Little Fishies Inc., Miami Gardens FL, USA). After two days’ acclimation, corals were brought to indoor flow-through seawater tanks where water temperatures could be controlled. Fragments from each genet were randomly allocated to either the ambient or high-temperature treatment, then randomly assigned to tanks within temperature treatments and rotated weekly within treatments to minimize tank effects. All fragments were illuminated to levels of photosynthetically active radiation similar to those on the reef (EcoTech Radion XR30w Pro, EcoTech Marine, Bethlehem PA, USA). Tanks were cleaned every two days to prevent algal overgrowth.

### Incubation

All tank temperatures were monitored and adjusted automatically at 15 min intervals for the duration of the experiment (APEX Controllers, Neptune Systems, Morgan Hill CA, USA). All tanks were programmed to include a diel fluctuation of 1°C. Ambient tanks ranged from 27° at night to 28° at midday so that daily averages remained well below the region’s coral bleaching threshold (28 + 1°C) throughout the experiment; this was also similar to diel flux in Kāne’ohe Bay from the preceding month [46]. For the heat stress treatment, the maximum daily tank temperature was increased by 1°C per day to a maximum of 31°C. After one week at a daily maximum of 31°C, an additional degree was added to the heat stress treatment so that the daily maximum was 32°C for one more week. After this two-week heated period, acute stress measurements were taken for both treatments and heated tanks were returned to ambient. The experiment continued for an additional 4.5 weeks during which the remaining corals were kept at ambient (~28°C) to investigate early recovery, after which measurements were taken again for the recovery time point (Fig. 1A). Accumulated experimental degree heating weeks (Fig. 1B) were calculated using tank daily mean temperatures [49].

**Figure 1.**
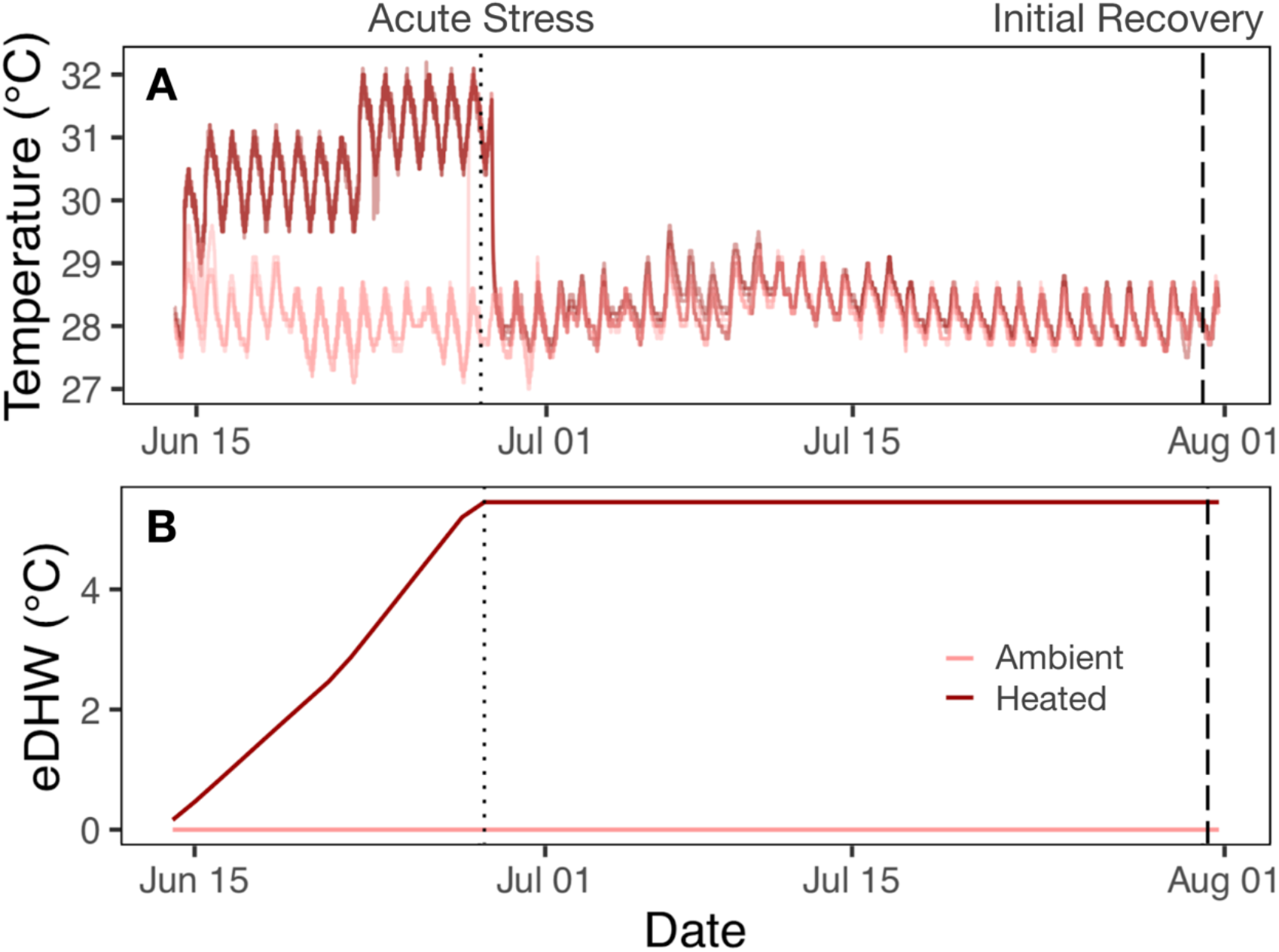
Experimental temperatures over stress and recovery periods. (A) Water temperatures in heated (N = 3 tanks) and ambient (N = 3 tanks) mesocosms, recorded in each tank every 15 minutes. (B) Accumulated daily experimental degree heating week (eDHW). Vertical lines indicate sampling timepoints: acute stress (dotted, June 28 2019) and initial recovery (dashed, July 31 2019).

### *In vivo* coral performance

Dark-adapted photochemical yield (F_v_/F_m_) was measured for each coral daily throughout the heat experimental period, and weekly throughout the initial recovery period. F_v_/F_m_ measurements were taken ~1 hr after sunset using a Diving-PAM (Walz GmbH, Effeltrich, Germany) 5-mm-diameter fiber-optic probe as described [46]. At the end of each experimental treatment period (acute stress and initial recovery), oxygen evolution of 20 fragments (N = 10 from each temperature history) was measured to calculate photosynthetic rate and light-enhanced dark respiration rate (LEDR) exactly as described [46] (Fig. S2). Fragments were chosen haphazardly so that each coral genet under both temperature regimes was represented.

### Stable isotope tracer experiment

After each respirometry experiment, those 10 fragments per temperature per timepoint (N = 40 fragments total) were placed upright on a rack in clear 10-gallon tubs of filtered seawater with pumps for circulation. 99% NaH^13^CO_3_ (Cambridge Isotope Labs, Tewksbury MA, USA) was added to non-limiting concentrations (31.2 μM; atom % enrichment NaH^13^CO_3_ = 2.678%). During this pulse, tubs were covered with plastic wrap to minimize ^13^CO_2_ offgassing and maintained at their experimental temperatures using an external water bath. After 7 hours in the light to incorporate the tracer, fragments were rinsed in a bucket of clean seawater and transferred to a darkened flow-through tank for a 12-hour chase period. Corals were then snap-frozen at −80°C until further processing.

### Physiological analyses

The 40 pulse-chase fragments were thawed on ice and airbrushed using filtered seawater (FSW) to remove all tissue. The resulting slurry was homogenized at 25,000 rpm for 10 s using a tissue homogenizer (Fisherbrand 850 Homogenizer, Fisher Scientific, Waltham MA, USA) and aliquoted for subsequent assays. Symbiodiniaceae concentrations were determined in triplicate within 6 hours of airbrushing using a flow-cytometer (Guava easyCyte 5HT, Luminex, Austin TX, USA) as described [46, 50]. If symbiont density could not be measured on D0, it was measured using a hemocytometer to avoid undercounting after chlorophyll deterioration. Chlorophyll was extracted in acetone, measured on a spectrophotometer (BioTek PowerWave XS2, Agilent, Santa Clara CA, USA), and concentration was calculated from the equations in Jeffrey & Humphrey 1965, all as described [46]. Protein concentration in the host fraction was measured on a spectrophotometer (BioTek ELx808, Agilent) using the Bradford method. Total host lipids were measured colorimetrically on a spectrophotometer (BioTek PowerWave XS2, Agilent) as described [51]. Surface area was determined by the single wax dipping method [52].

For ^13^C analysis, sample homogenates were separated by differential centrifugation into coral and algal fractions at 7000 x g for 5 min. The supernatant was removed as the host fraction, after which the pellet was washed in FSW and spun down again. The supernatant was again added to the host fraction. The symbiont fraction was then resuspended in FSW. Separated samples were then dried to stable weight (48 hrs at 50°C), enclosed tightly in tin capsules (EA Consumables, Marlton, NJ), and sent for ^13^C enrichment analysis by continuous-flow Isotope-Ratio Mass Spectrometry (UC-Davis Stable Isotope Facility, Davis CA, USA). A technical replicate was included for eight symbiont and host fractions.

### Measuring coral intracellular pH

For the initial recovery time point, one fragment per genet within each treatment was used to measure intracellular pH (pH_i_). Coral cells were isolated from each fragment and loaded with the pH-sensitive dye SNARF1-AM (Thermo Fisher Scientific) as described [46, 53]. Cells were pelleted, resuspended in filtered seawater, and imaged at 25°C in a glass-bottomed dish using an inverted confocal microscope (LSM 710, Zeiss, Oberkochen, Germany) with image acquisition settings identical to ref. [46]. At least eight gastrodermal cells containing algal symbionts (symbiocytes) were imaged from each coral. Corals lacking sufficient stained symbiocytes due to bleaching were excluded from the dataset. SNARF-1 fluorescence ratios within coral cytoplasm were quantified in ImageJ, normalized to background fluorescence, and converted to pH using a species-specific calibration curve generated on the same microscope, as described ([46]; Fig. S1).

### Statistical analysis

To test how corals responded to heat treatment, we generated linear mixed effects models for each physiological variable as a function of symbiont species (C and D) and treatment temperature history (28, 31), with genet as a random intercept (*lme4* package [54]). To avoid overfitting models given our population size, we generated separate models for both the acute and initial recovery experimental timepoints (Fig. 1, Table S2). To assess coral health over time, we modeled each physiological variable as a function of symbiont species and timepoint (acute stress and initial recovery), with genet as a random effect, generating separate models for both temperature histories (Fig. 1, Table S3). We examined each model’s Q-Q and residual plots to check that model residuals met assumptions of normality and constant variance. Where data were heavy-tailed, we used the *LambertW* package to Gaussianize the distribution. Significance of fixed effects and their interactions were determined using Satterthwaite’s Type III ANOVA. To test whether coral accumulation of symbiotic photosynthate affects host physiological function across treatments, we ran linear regressions (Pearson’s product-moment correlation) comparing host photosynthate accumulation with intracellular pH and calcification. To test how symbiont population dynamics and energy provisioning affected coral physiology, we modeled relationships between symbiont population traits and host physiology across treatments using either linear regressions or generalized additive models (where host response variables followed a non-monotonic distribution). All analyses were run in RStudio version 4.0.2 [55].

## RESULTS AND DISCUSSION

### Coral symbiont thermotolerance confers no nutritional benefit to hosts during heat treatment

Experimental heat treatment caused coral bleaching symptoms and colony stress regardless of symbiont species (Fig. 2). Symbiont photochemical efficiency (p < 0.0001), chlorophyll *α* concentration (p = 0.002), and light sensitivity of photosynthesis (*α*) (p = 0.011) all decreased for heat-treated corals relative to ambient (Fig. 2A, C, E), confirming that bleaching occurred. Although heat treatment did not significantly decrease symbiont cell density (p = 0.096, Fig. 2B), gross photosynthesis (p = 0.178, Fig. 2D), or fixed carbon incorporated into the symbiont fraction (^13^C at-%) (p = 0.095, Fig. 2F), heated corals assimilated less photosynthetically fixed carbon (p < 0.001; Fig. 2K). This signals a deficiency in photosynthate translocation to the coral host after heat stress, consistent with previous empirical [42–44] and theoretical [56] studies. Possibly as a result, heat-treated coral colonies also calcified less (p = 0.044, Fig. 2J) (Fig. 2B). However, regardless of heat treatment, corals hosting *Cladocopium* sp. algae (hereafter C-colonies) incorporated more fixed carbon overall than corals hosting *Durusdinium glynni* (hereafter D-colonies) (p = 0.018; Fig. 2K). This contradicted our hypothesis, and was surprising given that carbon benefits to C-corals can be lost at higher temperatures in other host species [30]. Our results instead suggest that *M. capitata* hosting *Cladocopium* sp. retain a photosynthate incorporation advantage over D-colonies during heat stress.

**Figure 2:**
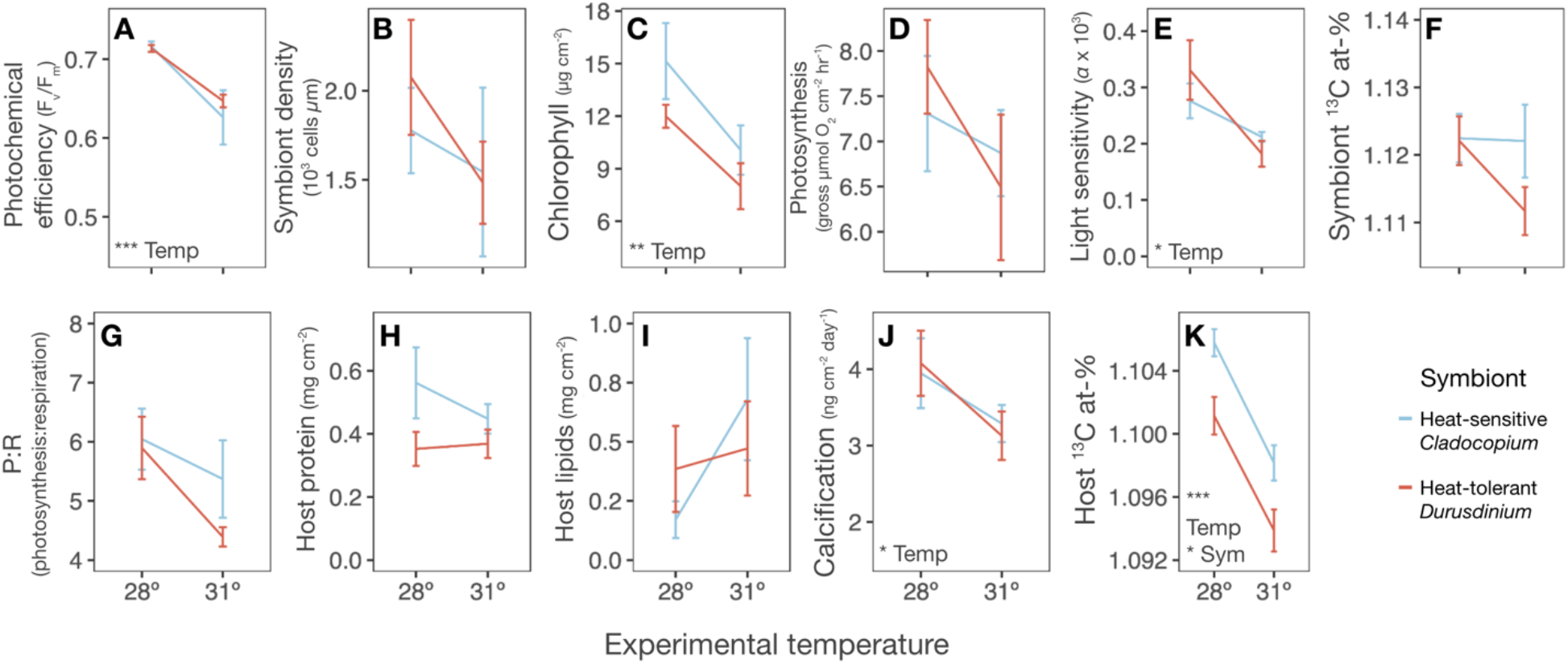
Experimental heating caused bleaching and physiological stress in *Montipora capitata* regardless of symbiont association. **A-F)** Heat decreased symbiont photochemical efficiency (F_v_/F_m_) **(A)**, chlorophyll **(C)**, and light sensitivity (*a*) **(E)** though it did not affect symbiont areal density **(B)**, gross photosynthesis **(D)**, or net carbon fixation **(F)**. **G-K)** Heat decreased coral calcification **(J)** and net fixed carbon assimilation (^13^C atom %) **(K)** although it did not affect photosynthesis:respiration **(G)**, or total host protein **(H)** or lipids **(I)**. Coral photosynthate assimilation (host ^13^C atom-%) also was higher for C-corals than for D-corals regardless of temperature **(K)**. Error bars = SEM. N = 5 genets per group. Insets show results of linear mixed effects models with temperature (Temp) and symbiont (Sym), with coral genet as a random intercept (* = p < 0.05, ** = p < 0.01, *** = p < 0.001). Full model results are reported in Table S2.

### Health of host and symbiont population are uncoupled during initial bleaching recovery

One month following cessation of heat stress, *Durusdinium glynni* maintained steady photochemical efficiency and began recovering to higher population density within the host, while *Cladocopium* sp. continued to lose photochemical efficiency and decline in population density within the host (F_v_/F_m_ ~ Timepoint * Symbiont, p = 0.046; Symbiont Density ~ Timepoint * Symbiont, p = 0.034; Fig. 3A-B). This recovery in heat-stressed D-colonies led to comparable symbiont density and chlorophyll content relative to ambient colonies at the end of the experiment, while heat-stressed *Cladocopium-*dominated colonies still had fewer symbionts (Temperature * Symbiont, p = 0.014) and less chlorophyll (Temperature * Symbiont, p = 0.008) than ambient colonies (Fig. S3B-C). Surprisingly, this early recovery of *D. glynni* did not give D-colony hosts any significant advantage over C-colony hosts in terms of P:R, protein, lipids, calcification, or host carbon assimilation (p ≥ 0.210: Fig. 3G-K, Fig. S3G-K). Thus, though *D. glynni* symbionts recovered faster than *Cladocopium* sp., neither host group showed signs of recovery within 4 weeks of the heat stress. This contradicts our hypothesis that the faster-recovering *D. glynni* [47] offers corals an advantage after heat stress. However, delayed host recovery is consistent with observations that endosymbiont populations rebound faster than the host during initial bleaching recovery [57–59]. Host lipids continued to deplete after heat stress abated (p = 0.027, Fig. 3I), which is consistent with prior evidence that corals remain energetically limited during bleaching recovery [60], and in particular that this population of *M. capitata* catabolizes lipids to survive bleaching [61] and takes at least 6 weeks to replenish energy reserves [62]. Given that *M. capitata* can also compensate for lost symbionts by increasing heterotrophy [63], the sluggish early recovery we observed may have been mitigated if we had fed corals during the experiment. Furthermore, even ambient-treated corals experienced physiological stress in our mesocosms, displaying lower photochemical efficiency (p < 0.0001, Fig. S4A), chlorophyll concentration (p = 0.002, Fig. S4C), calcification rates (p < 0.0001, Fig. S4J), and host carbon assimilation (p < 0.001, Fig. S4K) over time. This may have been due to sub-bleaching heat stress *in situ* prior to sampling and/or during the experiment: Kane’ohe Bay was already anomalously warm by the time of collection in June 2019 [46, 47] and ambient tank temperatures peaked above 29°C (these corals’ projected bleaching threshold) several times during the experiment (Fig. 1A). However, all corals were exposed equally to these additional stressors, and they do not alter our result that faster-recovering *D. glynni* symbionts were not sufficient to improve *M. capitata* carbon assimilation, biomass, or skeletal growth rates after bleaching.

**Figure 3.**
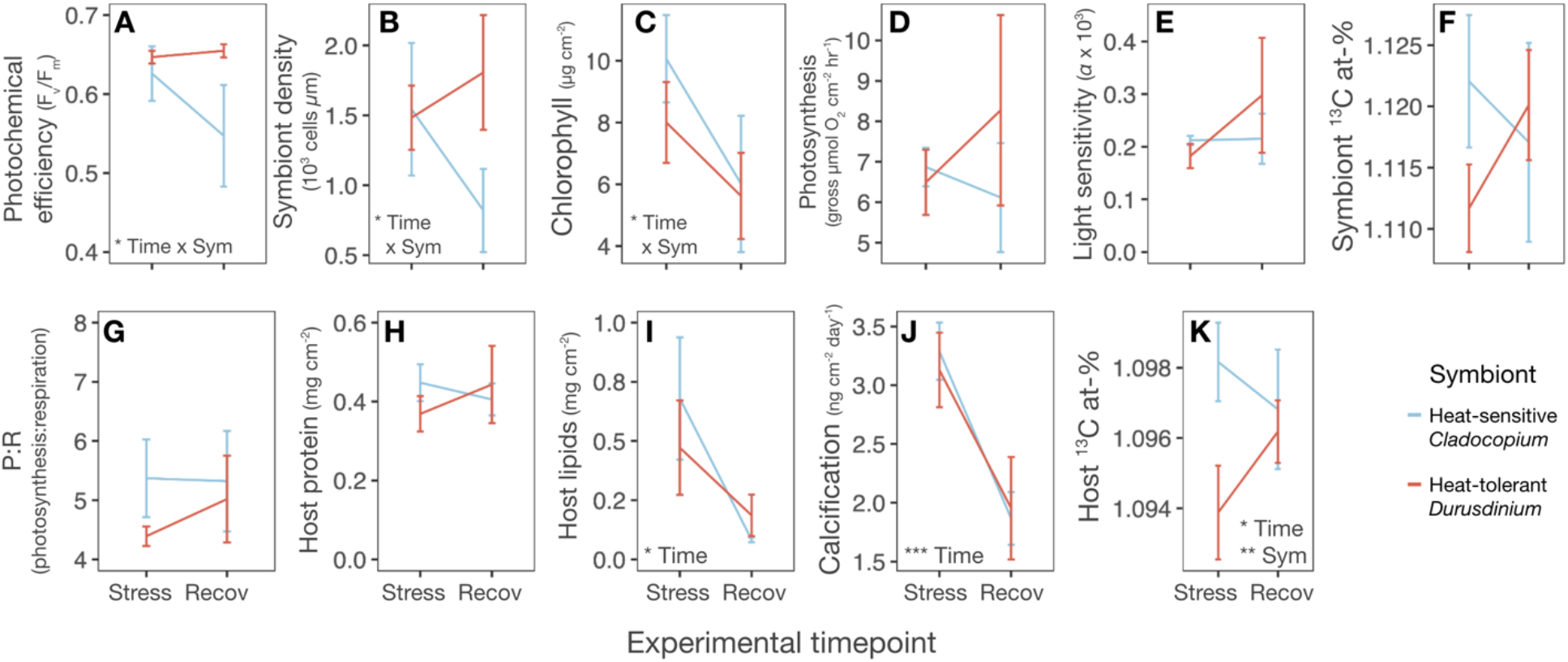
Symbiont and host physiology trajectories are uncoupled during the month following heat stress. **A-F)** Thermotolerant and thermosensitive symbionts responded differently after heat stress: thermotolerant *Durusdinium* retained or improved photochemical efficiency (F_v_/F_m_) **(A)** and symbiont density **(B)** over initial recovery (‘Recov’) after acute stress (‘Stress’), whereas thermosensitive *Cladocopium* continued to lose both. *Cladocopium* also lost more chlorophyll **(C)** than *Durusdinium* did. Gross photosynthesis **(D)**, light sensitivity of photosynthesis **(E)**, and symbiont carbon fixation **(F)** were unaffected by timepoint or symbiont species. **G-K)** *Durusdinium* thermotolerance did not improve host physiology following heat stress: symbiont species did not affect photosynthesis:respiration **(G)**, total host protein **(H)** or lipids **(I)**, or calcification **(J)**, and C-corals assimilated more photosynthate overall (host ^13^C at-%) **(K)**. Error bars = SEM. N = 5 genets per group. Insets show results of linear mixed effects models with timepoint (Time) and symbiont (Symb), with coral genet as a random intercept (* = p < 0.05, ** = p < 0.01, *** = p < 0.001). Full model results reported in Table S3.

### Coral intracellular pH depends on symbiotic benefits

The effect of heat stress on coral intracellular pH (pH_i_) was dependent on symbiont species. Specifically, prior exposure to heat stress led to decreased pH_i_ in recovering D-colonies (p = 0.018); however, C-colonies resisted this acidification and maintained pH_i_ in the aftermath of bleaching (Temperature * Symbiont, p = 0.036; Fig. 4A). Since D-colonies tend to be more bleaching-resistant, and coral species bleaching susceptibility predicts the degree of pH_i_ dysregulation after heat stress [64], we had expected D-colonies’ pH_i_ to be more resilient to heat treatment. C-colonies in this study may have had a pH_i_ regulatory advantage over D-colonies because *Cladocopium* sp. continued translocating more fixed carbon than *D. glynni* to their hosts despite acute heat stress (Fig. 2K), providing the host with the energy needed to maintain acid-base homeostasis. Interestingly, the greater acidification we observed here in D-colonies inverts the pattern observed for these same corals during a marine heatwave *in situ*, when C-colonies had more acidic pH_i_ than D-colonies, attributed to their greater bleaching susceptibility [46]. The advantage *M. capitata* obtains from hosting certain symbionts thus likely depends on the duration and severity of heat stress, as well as the timing of measurement relative to stress, since the *in situ* study measured pH_i_ at peak heat stress rather than after 4 weeks of recovery, as was done here.

**Figure 4:**
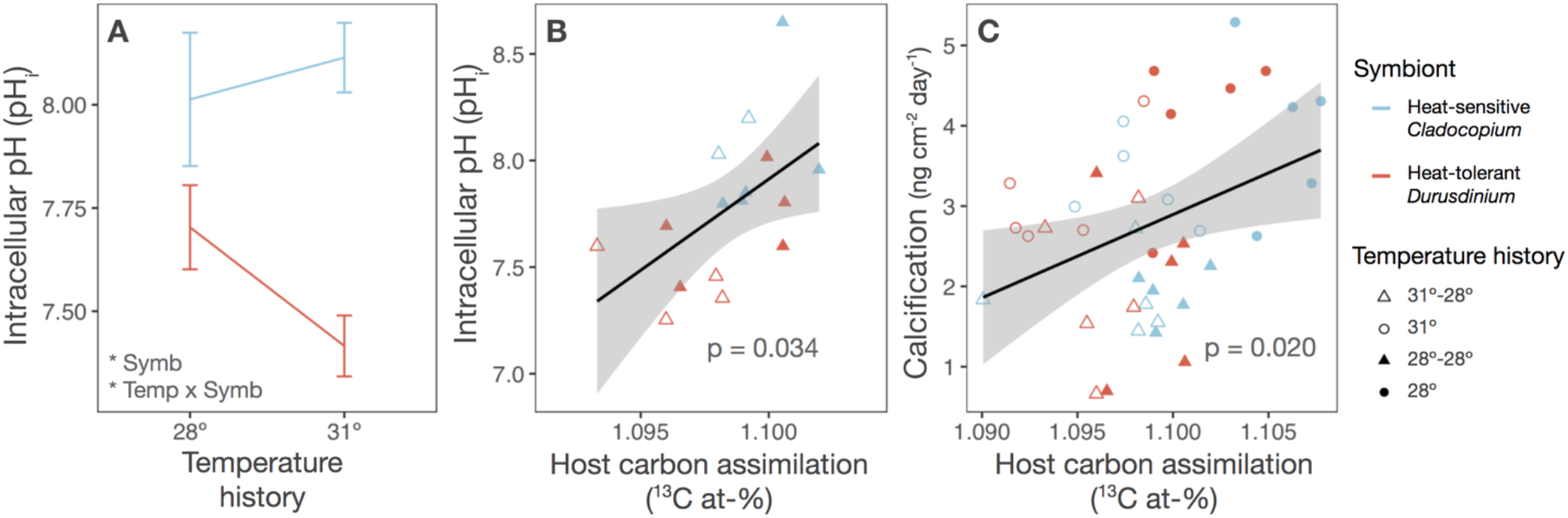
Assimilated carbon from symbiosis predicts coral calcification and intracellular pH after heat stress. **A)** Symbiont thermotolerance affects how temperature alters coral symbiocyte intracellular pH (pH_i_) during bleaching recovery. Inset shows result of linear mixed effects model with symbiont species (Symb) and temperature treatment (Temp), with coral genet as a random intercept (* = p < 0.05). Error bars = SEM. Full model results reported in Table S2. **B)** Host fixed carbon assimilation is positively correlated with symbiocyte pH_i_. Gray line, linear correlation; inset, linear regression p-value (t = 2.3506, df = 14, R^2^ = 0.2830). **C)** Host fixed carbon assimilation predicts colony calcification rates. inset, linear correlation p-value (t = 2.438, df = 35, R^2^ = 0.1452). Black lines show linear regressions; gray area = 95% confidence interval.

To understand how variable carbon translocation may have impacted coral cell physiology, we explored correlations between coral pH_i_ and host carbon assimilation and found that host carbon assimilation predicted intracellular pH (pH_i_) across experimental groups (R^2^ = 0.283, p = 0.034; Fig 3B). This result suggests corals deriving more energy from symbiosis are better equipped to maintain acid-base homeostasis, consistent with the explanation that D-colonies had lower overall pH_i_ than C-colonies because their algae translocated less photosynthate (Fig. 2K). Alternatively, pH_i_ differences may have resulted from variation in endosymbiont metabolic rates, as dinoflagellate photosynthesis increases coral cell pH_i_ via CO_2_ drawdown [53, 65, 66]. However, corals in this experiment were dark-adapted prior to measuring pH_i_, likely minimizing any residual differences in pCO_2_ and pH_i_ due to algal photosynthesis. Indeed, simple linear regression analysis confirmed that there was no correlation between per-alga photosynthetic rate and host pH_i_ (Fig. S5). Therefore the cellular acidification in corals hosting *Durusdinium* was more likely a result of the organismal-level deficiencies in energy provisioning we observed in D-colonies.

### Potential pH regulatory tradeoffs

Similarly to pH_i_, colony calcification rate was directly related to coral host photosynthate assimilation (p = 0.020, R^2^ = 0.134, Fig. 4C), suggesting that corals receiving more carbon from their symbionts can maintain higher skeletal growth rates. However, despite the parallel correlation between symbiocyte pH_i_ and host photosynthate assimilation (Fig. 4B), colony calcification was not related to pH_i_ (p = 0.792, R^2^ = 0.005, Fig. S6). Because corals use some similar mechanisms to control pH_i_ [53] as well as pH of the extracellular calcifying medium (pH_ECM_) [67], we had expected corals with acidified pH_i_ to be less able to maintain alkaline pH_ECM_ and thus calcification. But coral pH regulation may be subject to tradeoffs between homeostasis and growth. Since these corals were early in recovery [60, 62] and still showing signs of accumulated metabolic stress (Fig. 3I-J), they likely had limited available resources to invest in pH regulation to simultaneously maintain pH_i_ and calcification. Corals might thus prioritize maintaining pH_i_ at the expense of pH_ECM_ during bleaching recovery, perhaps by altering protein expression or localization of different ion transporters [68], obscuring any individual-level differences in total pH regulatory capacity. This sort of ‘preferential pH regulation’ has been observed in air-breathing fishes that rapidly regulate pH_i_ in response to CO_2_ stress while allowing extracellular pH to remain acidic [69, 70]. Further research into the relationship between coral pH_i_ and calcification should test the possibility that cnidarians also use preferential pH_i_ regulation to balance the multiple acid-base challenges they face, including endosymbiont photosynthesis, cellular respiration, environmental acidification, and calcification [39].

### Symbiont strategic shift toward parasitism

Surprisingly, algal density within host tissue failed to predict host physiology across Symbiodiniaceae species and temperature treatments. We originally hypothesized that corals retaining more symbionts during and after heat stress would assimilate more photosynthate. Because symbiont photosynthesis provides a majority of these corals’ energy [31], we tested for correlations between symbiont cell density and host performance. Higher algal population density did not correlate with greater host carbon assimilation (R^2^ = 0.030, p = 0.289; Fig. 5A), lipid content (p = 0.292; Fig. S8A) or calcification rate (R^2^ = 0.088, p = 0.064; Fig. S8B). Instead, Symbiodiniaceae cell density was directly correlated with the amount of labeled carbon that symbionts retained instead of translocating to the host (p = 0.017, R^2^ = 0.141; Fig. 5B), a measure of symbiont ‘selfishness’ [42]. We obtained a similar result calculating algal cell density areally (p = 0.012, Fig. S7A), confirming that the effect was not an artifact of differences in host protein content [71, 72] and that a symbiont population’s selfishness was directly related to its density.

**Figure 5.**
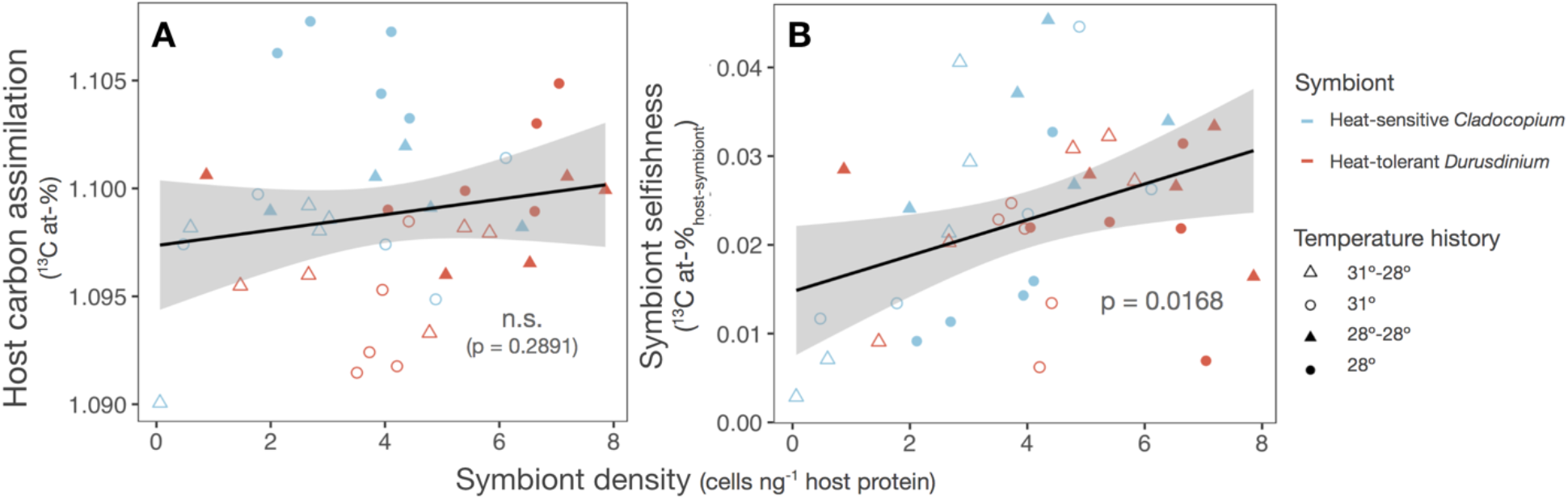
During and after heat stress, symbiont population growth increases selfishness. **A)** Symbiont density in host tissue does not predict host fixed carbon accumulation. Inset shows result of linear regression (t = 1.075, df = 38, R^2^ = 0.0295). **B)** Symbiont density is positively correlated with selfishness. Selfishness was calculated by subtracting ^13^C enrichment (^13^C at-%) in the host fraction from ^13^C at-% in the symbiont. Inset shows result of linear regression (t = 2.5018, df = 38, R^2^ = 0.1414). Black lines show linear regressions; gray areas = 95% confidence intervals.

One plausible explanation is that high-density symbiont populations translocated less of their fixed carbon in response to symbiont self-shading [40, 41, 73]. However, we saw no signs of self-shading in our study. Light-limited symbionts typically upregulate chlorophyll to compensate [74, 75], but we instead found that per-symbiont chlorophyll was lowest at high symbiont densities (p < 0.0001; Fig. S8C). Furthermore, colony gross photosynthesis was not sufficient to predict coral carbon assimilation (R^2^ = 0.003, p = 0.743, Fig. S9), confirming that differences in total symbiont productivity could not account for symbiont cell density increasing selfishness. Another alternative explanation is that corals hosting more symbionts incurred greater respiration costs during the pulse-chase experiment, resulting in lower host carbon assimilation. In fact, total colony respiration did increase with symbiont density (R^2^ = 0.107, p = 0.042; Fig. S8D), confirming that high symbiont concentrations increase costs to the host (e.g. [40, 41, 72, 73]; but see [76]). However, we found no correlation between total colony respiration and calculated symbiont selfishness (R^2^ = 0.012, p = 0.517, Fig. S7B), indicating the density-selfishness relationship is not simply because increased respiratory burden decreases host carbon assimilation. Because neither algal primary productivity nor colony respiration costs explain calculated symbiont selfishness, high-density symbionts must be sequestering a greater proportion of the carbon they fix.

Our finding that multiple species of Symbiodiniaceae retain more photosynthate at high cell densities (Fig. 5B) suggests the symbionts remaining *in hospite* after coral bleaching can alter their nutrient allocation strategy in response to changes in density. If Symbiodiniaceae divert photosynthate from cooperating with the coral to their own cell division, achieving high algal population density may require a corresponding increase in selfishness (Fig. 6). At ambient conditions, corals invest ATP in regulating inorganic nitrogen and carbon inside the symbiosome [77, 78], while the symbiont translocates much of its resulting photosynthate back to the coral [13] (Fig. 6A). After bleaching, however, algae may prioritize their recovery by translocating less fixed carbon to the coral host (Fig. 2K; Fig. 5B; Fig. 6B) even as a larger algal population incurs additional costs to the coral host [40, 41], further limiting the host’s available energy (Fig. 6B). Such a strategic shift at the expense of the coral could explain why energetic losses from bleaching can persist for months (e.g. [60]), and why sexual reproduction can suffer even years after colonies regain normal symbiont populations (e.g. [79, 80]).

**6:**
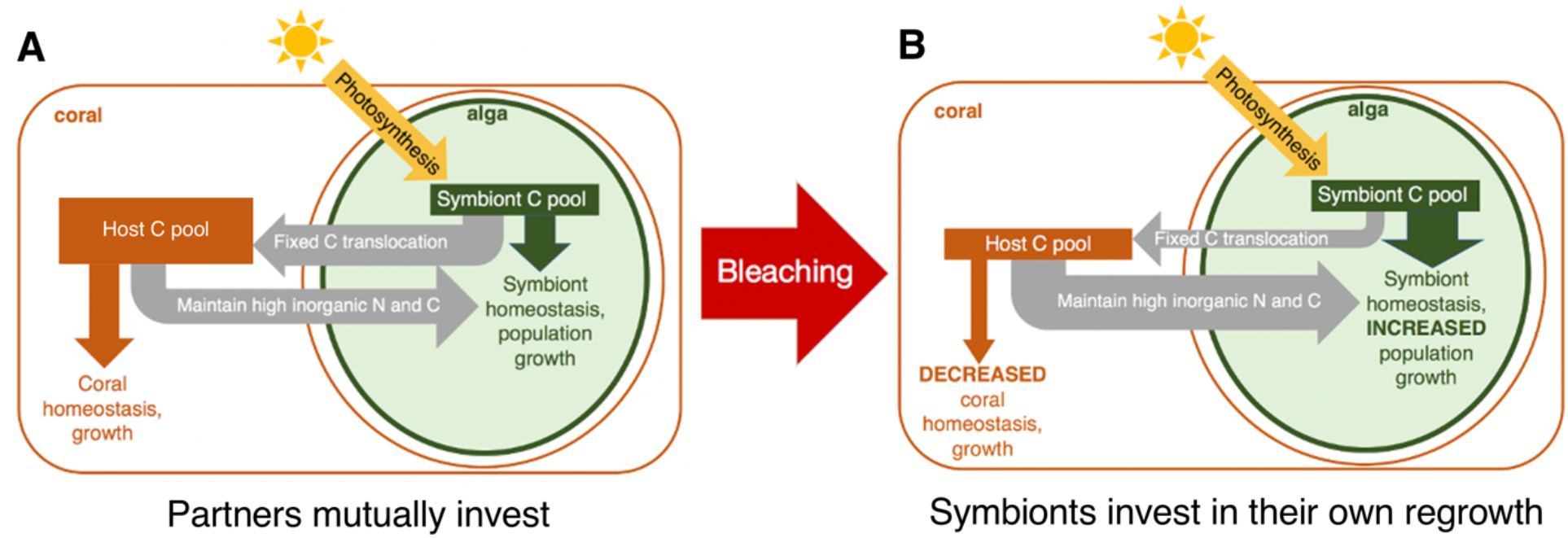
Model of symbiont strategic shift. **A)** Without stress, coral investment in symbiont photosynthesis and growth (lower gray arrow) produces returns in photosynthetically fixed carbon translocated to the coral (top gray arrow). **B)** After a bleaching event reduces symbiont population density, symbionts may invest in their own cell division and regrowth (larger green arrow) at the expense of translocating fixed carbon to the coral (smaller gray arrow). The coral is left with fewer resources (smaller orange box) for its own growth and recovery (smaller orange arrow) while maintaining a growing symbiont population (larger gray arrow).

While some symbionts are known to behave more parasitically at high densities (e.g. in the Branchiobdellid-crayfish cleaning symbiosis [81]), this study is to our knowledge the first evidence suggesting a nutritional endosymbiont could maintain or achieve higher densities during stress by cooperating less with its host. It may also help explain observations that corals benefit most from intermediate symbiont densities [40, 41, 72]. Most existing carbon-acquisition-focused frameworks of coral bleaching obtain from intrinsic density dependence (i.e. symbionts becoming light- and CO_2_-limited) and increased respiratory costs of high symbiont loads; they assume each symbiont population translocates a constant proportion of their available photosynthate. Our results instead add to the evidence that corals’ optimal symbiont abundance also depends on how the algae allocate their fixed carbon (e.g. [43, 56]). Therefore, symbiont nutrient exchange strategy may itself be a dynamic regulator of bleaching susceptibility. High symbiont cell densities can increase bleaching risk (e.g. [71, 82]; but see [32]), a phenomenon thought to result from enhanced accumulation of reactive oxygen species as a result of symbiont photostress (e.g. [83–85]). However, heat-induced bleaching in cnidarians can occur without increases in reactive oxygen [86], signatures of oxidative damage [87], or even photostress [88]. The positive relationship we find between symbiont cell density and selfishness provides an alternative mechanism by which bleaching likelihood could depend on symbiont density. Diminished but fast-regrowing symbiont populations may leave corals carbon-deprived, poorly equipped to face the energetic demands of heat stress, and thus more vulnerable to the carbon limitation theorized to cause bleaching [43, 56, 73]. Future bleaching recovery research should correlate symbiont *in vivo* regrowth rates with carbon translocation to better validate this framework.

### Implications for coral survival under repeated heatwaves

Though immediately costly to the coral, subsidizing algal regrowth could be a viable strategy for long-term fitness: the faster a host regains equilibrium symbiont density, the sooner it will restore its full autotrophic potential. However, with some reefs predicted to bleach annually by 2043 under the current emissions scenario [89], even corals that recover all their symbionts quickly after bleaching (e.g. [23, 62]) will no longer be able to sustain stressful cycles of algal regrowth, complicating attempts to predict which corals could be ‘climate-proof’ [39, 90, 91][92, 93], [39, 90, 91]. It is therefore crucial to prioritize resilient and sustainable mutualism function in research aiming to support coral conservation. Assisted evolution *in vitro* to improve Symbiodiniaceae intrinsic thermotolerance (e.g. [94]) or colonization (e.g. [95]) is a promising new approach, and future work should test that the experimentally evolved algae still provide enough nutritional benefits to the coral host. Indeed, artificial selection for endosymbiont mutualistic traits can endanger mutualism stability when selection is performed outside of the host [96]. As the most thermotolerant coral holobionts appear to rely on extensive coral-algal coevolution rather than symbiont shuffling [97], future coral bleaching resilience studies must go beyond measuring endosymbiont growth and instead measure partner traits that favor functional stability of this critical mutualism [98].

More broadly, predicting mutualisms’ resilience to climate change requires understanding the actual function of the interaction under stress. We found that deficiencies in symbiotic nutrient exchange uncouple partner recovery trajectories during mutualism reestablishment. Researchers must therefore distinguish between the separate processes of symbiont regrowth and mutualism recovery when one or more partners can withhold resources to favor reproduction over cooperation. Finally, reestablishing a beneficial endosymbiosis can itself pose a secondary challenge for holobionts surviving stress, highlighting the need for more mechanistic study of how mutualisms recover from stress in our current era of unprecedented environmental change.

## DATA AVAILABILITY

Data and R code are available on GitHub at github.com/allenwaller/MontiporaSymbStrategy

## ACKNOWLEDGEMENTS

We thank Teegan Innis, Crawford Drury, Josh Hancock, Ariana Huffmyer, Shayle Matsuda, Mindy Mizobe, and the staff of the Hawai’i Institute of Marine Biology for logistical support during the experiment; Christopher Carlson for image analysis assistance; Christopher Wall for indispensable isotope advice; and the staff of the UC-Davis Stable Isotope Lab for isotope analysis. Corals were collected under Special Activity Permit 2020-41. This work was supported by the NIH T32 Predoctoral Training Grant in Cell and Molecular Biology GM-07229 to L.A.-W., NSF-OCE 1923743 to K.L.B, Charles E. Kaufman Foundation New Investigator Award to K.L.B., and funding from the University of Pennsylvania to K.L.B.

