## Supplemental Figures & Table 1 for "Endosymbiont strategic shifts inhibit cooperation during coral bleaching recovery"

**Table S1.** Colonies sampled for this experiment. Symbiont percentages reflect qPCR results from colony samples taken July 9, 2019, reported in Dilworth et al. 2020. SD = standard deviation of the mean.

| Colony | % *Durusdinium* | % *Durusdinium* SD | Bleaching history |
|---|---|---|---|
| 11 | 0 | 0 | Susceptible |
| 19 | 0 | 0 | Susceptible |
| 201 | 0 | 0 | Susceptible |
| 203 | 0 | 0 | Susceptible |
| 211 | 0 | 0 | Susceptible |
| 202 | 82.8 | 25.16 | Resistant |
| 222 | 84.8 | 17.21 | Resistant |
| 20 | 95 | 4.690 | Resistant |
| 12 | 97.8 | 1.483 | Resistant |
| 214 | 98.6 | 1.517 | Resistant |

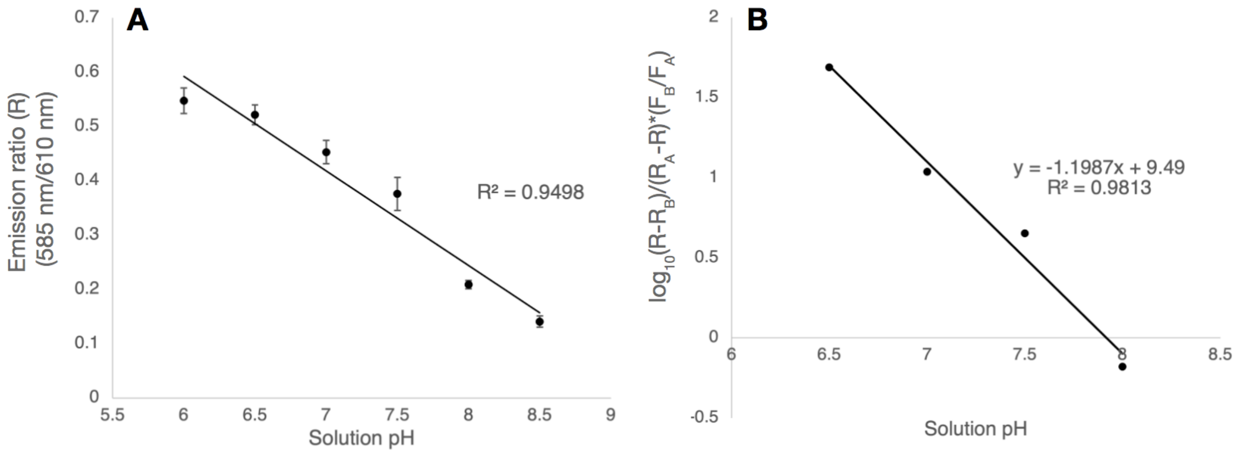


**Figure S1.** *In vivo* calibration of the pH-sensitive dye SNARF1 in *Montipora capitata* cells. A) Direct relationship between calibration solution pH and ratio (R) of SNARF emission at 585 ± 10 nm to 610 ± 10 nm. B) R was related to pH using the following equation: $pH = pK_A - \log_{10}(R-RB)(RA-R)*(FB/FA)$ where $R_A$ = emission ratio at pH 8.5, $R_B$ = emission ratio at 6, $F_B$ = 610 nm emission at pH 8.5, $F_A$ = 610 nm emission at pH 6, and $pK_A$ = x-intercept obtained from plotting the standard's logarithmic term against solution pH (shown).

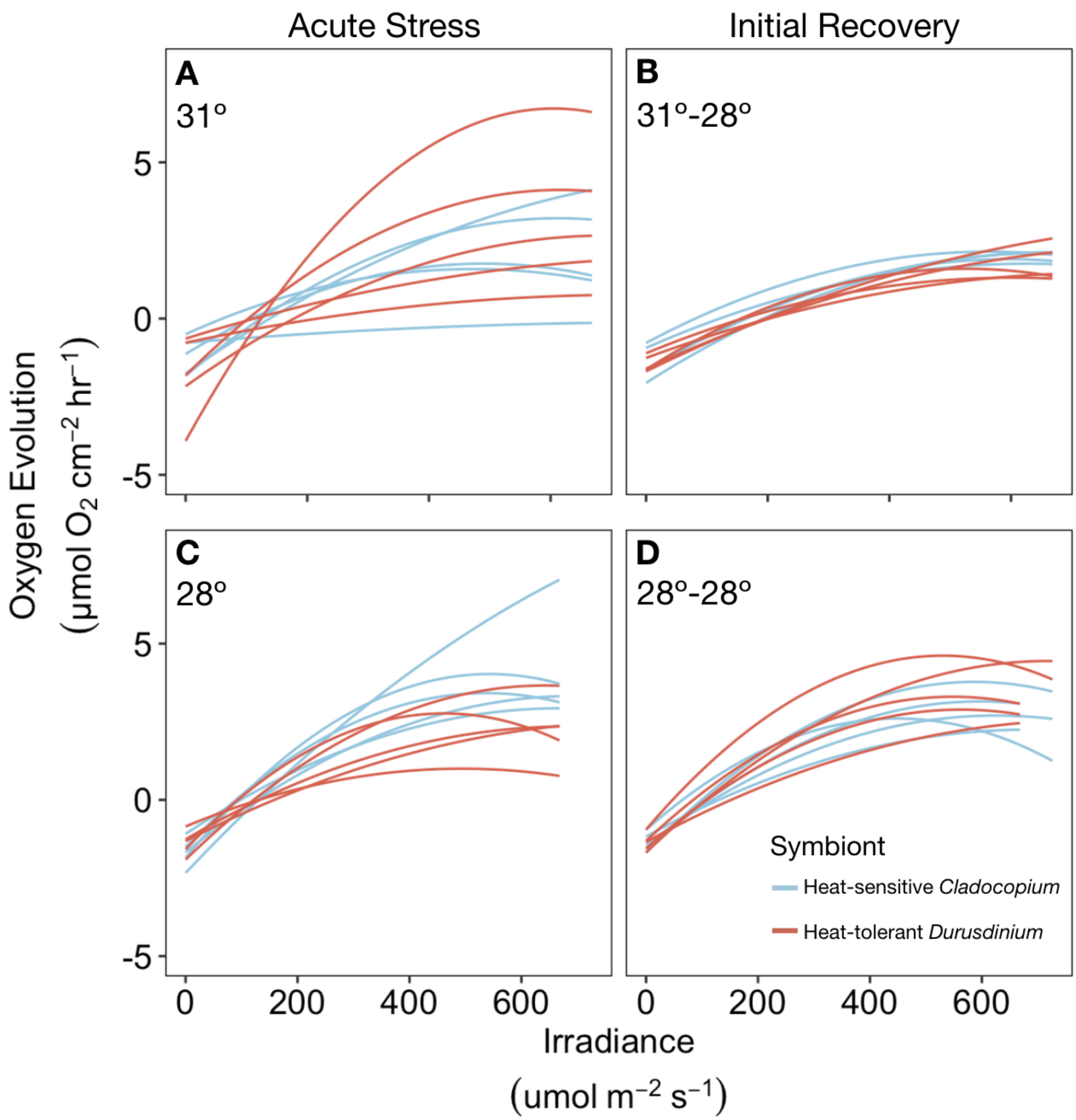


**Figure S2.** Photosynthesis-irradiance curves of heat-treated (A-B) and control temperature (C-D) *M. capitata* generated by curve fitting of oxygen consumption and production rates at the acute stress (A, C) and initial recovery (B, D) timepoints.

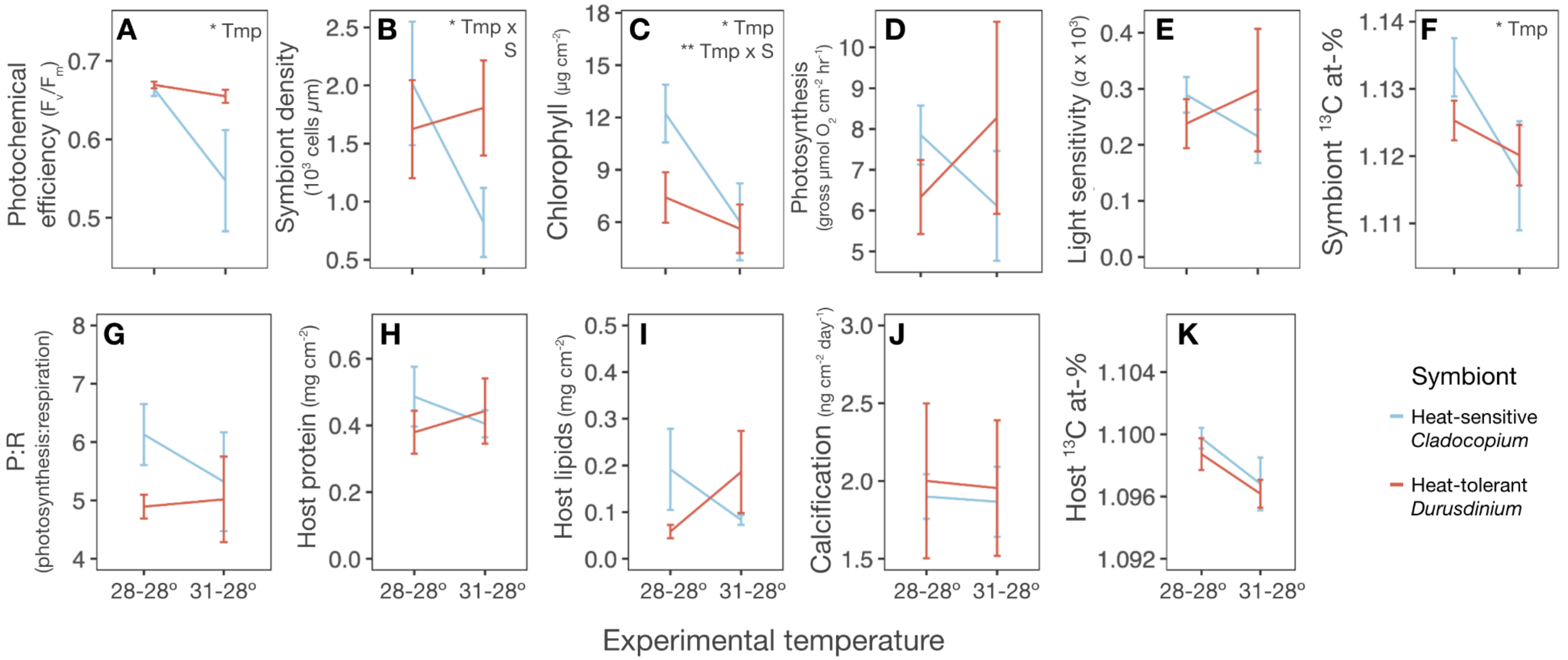


**Figure S3.** Endosymbiont and host responses after initial recovery from heat stress. **A-F)** Symbiont populations with a history of heat stress lost photochemical efficiency (Fv/Fm) **(A)** and fixed less carbon (symbiont $^{13}C$ atom %) **(F)** regardless of symbiont species, though gross photosynthesis **(D)** and light sensitivity of photosynthesis **(E)** were unaffected. Populations of *Cladocopium* sp. also lost more symbiont cell density **(B)** and chlorophyll **(B)** after heat stress than populations of *Durusdinium glynni*. **G-K)** Neither symbiont type nor temperature history significantly affected host physiology one month after heat stress. Insets show results of linear mixed effects models with temperature (Tmp) and symbiont (S), with coral genet as a random intercept (* = $p < 0.05$, ** = $p < 0.01$, *** = $p < 0.001$). Error bars = SEM. N = 5 genets per group. Full model results reported in Table S2.

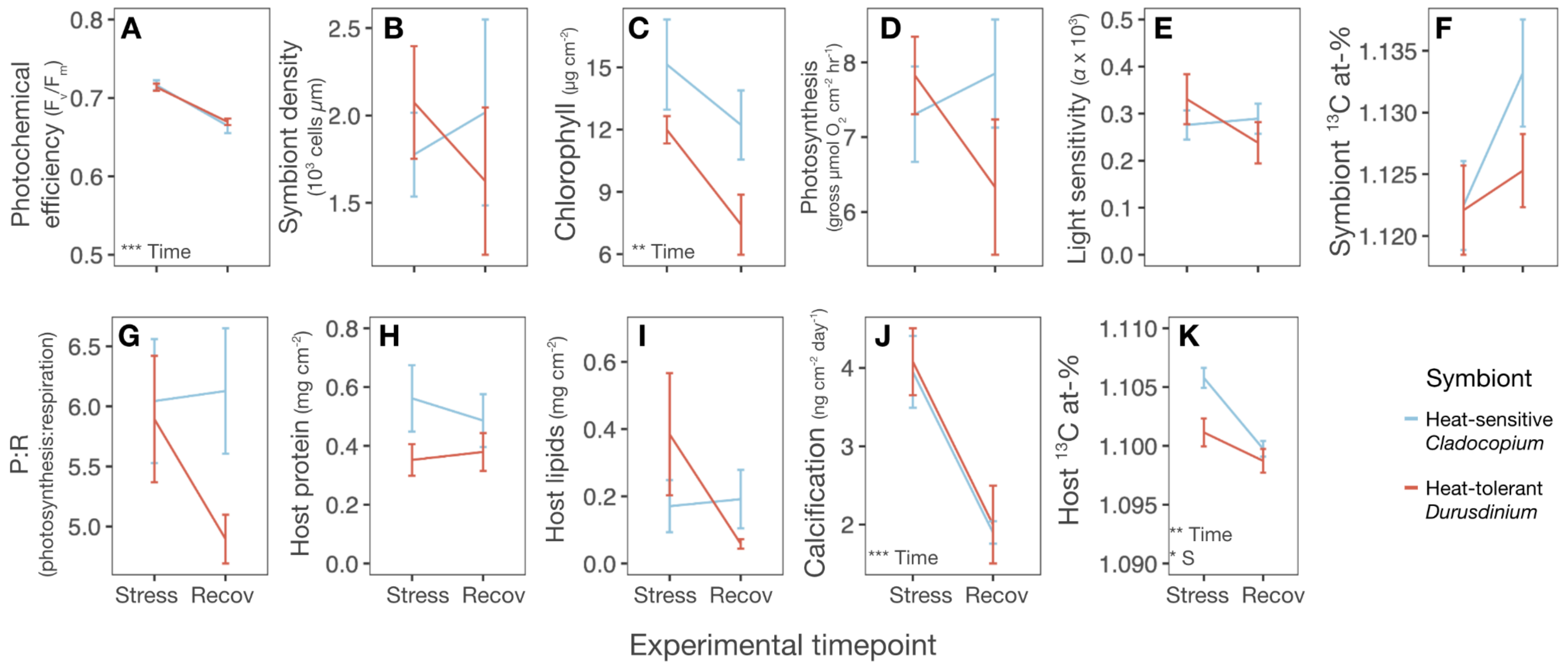


**Figure S4.** Trajectories of ambient-temperature *M. capitata* over time. A-F) Ambient-treatment symbiont populations lost photosynthetic efficiency (A) and chlorophyll (C) over the experimental time period but did not change in cell density (B), gross photosynthesis (D), light sensitivity (E), or carbon fixation (F). G-K) Ambient-treated corals did not change in photosynthesis:respiration ratio (G), total protein (H), or lipid (I) density, but they slowed calcification (J) and assimilated less fixed carbon from photosynthesis ($^{13}$C atom-%) (K) over the experiment. D-corals assimilated less photosynthate overall (K). This figure represents ambient-temperature fragments only, so x-axis labels refer to timepoint corresponding with Fig. 2: Stress = acute stress timepoint; Recov = initial recovery timepoint. Insets show results of linear mixed effects models with timepoint (Time) and symbiont species (S), with coral individual as a random intercept (* = $p < 0.05$, ** = $p < 0.01$, *** = $p < 0.001$). Error bars = SEM. N = 5 genets per group. Full model results reported in Table S3.

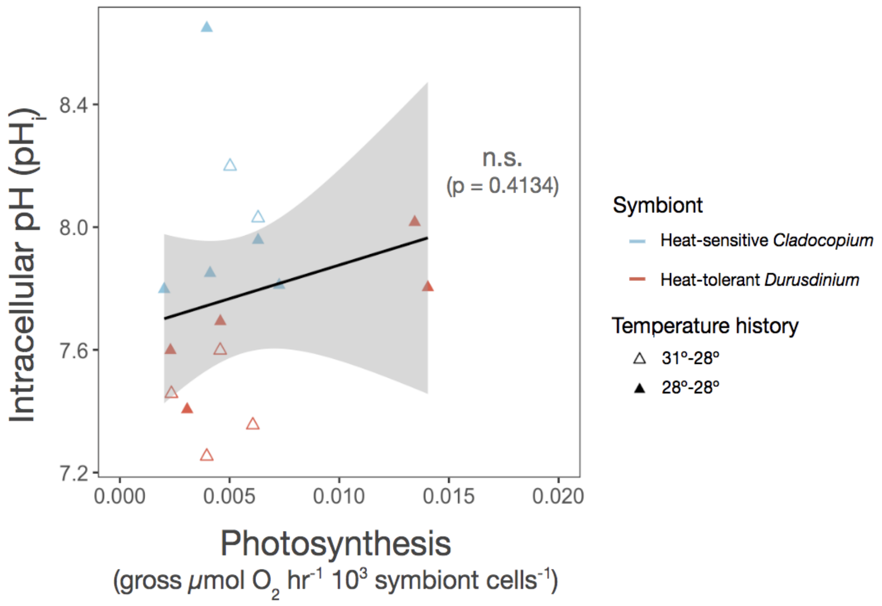


**Figure S5.** Symbiont photosynthesis does not increase *M. capitata* symbiocyte intracellular pH ($pH_i$) after dark-acclimation ($t = 0.84295$, $df = 14$, $R^2 = 0.0483$, $p = 0.4134$).

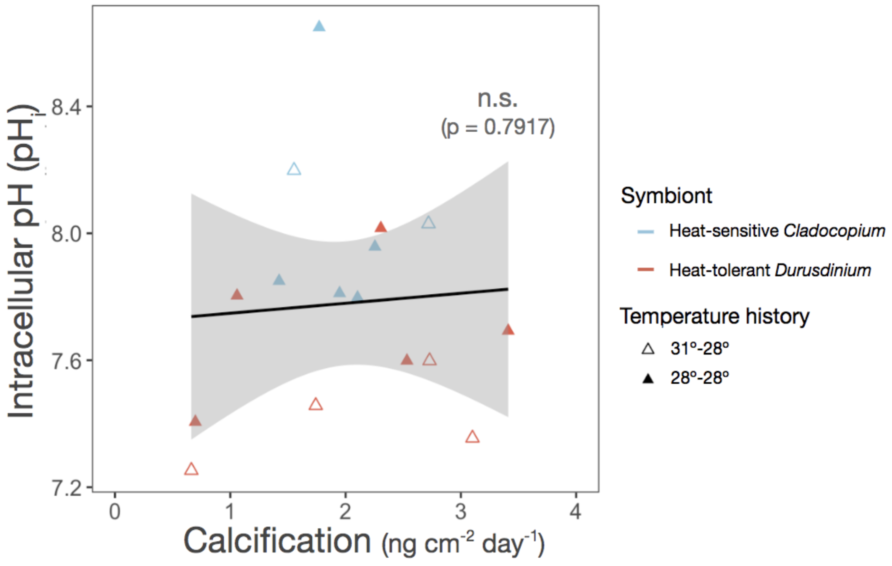


**Figure S6.** *M. capitata* symbiocyte intracellular pH ($pH_i$) does not predict colony calcification rate ($t = 0.26914$, $df = 14$, $R^2 = 0.0051$, $p = 0.7917$).

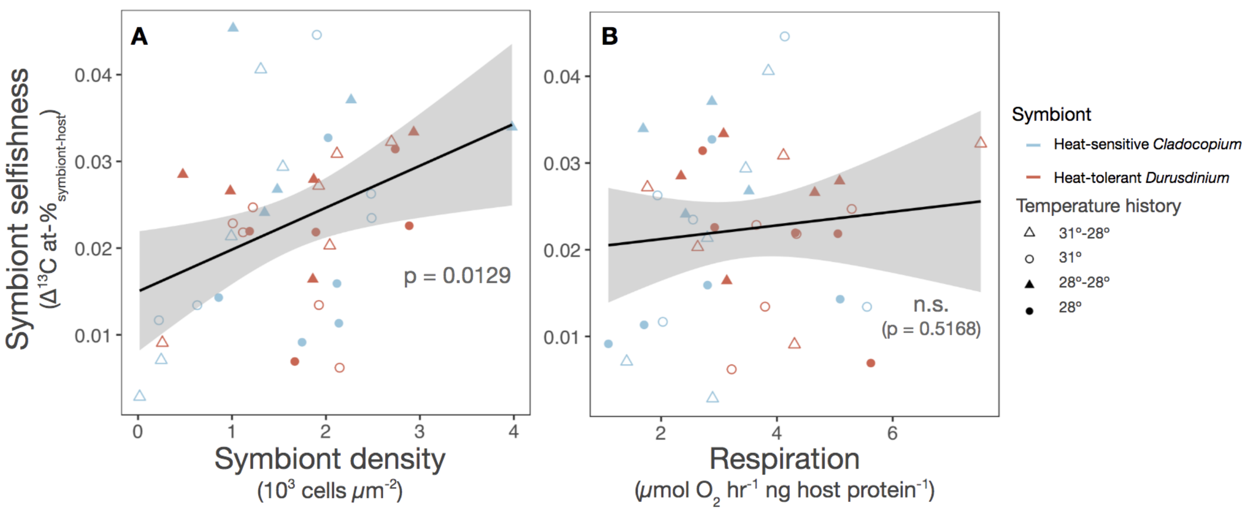


**Figure S7.** Symbiont selfishness ($\Delta^{13}C$ atom-$\%_{symbiont\text{-}host}$) increases with symbiont density independent of colony respiration. A) Areal symbiont density is positively correlated with the proportion of photosynthate symbionts retained instead of translocating to the coral host (t = 2.6087, df = 38, $R^2$ = 0.15188, p = 0.01292). Full linear mixed effects model results reported in Table S4. B) Colony respiration does not predict symbiont photosynthate retention (t = 0.65453, df = 37, $R^2$ = 0.01145, p = 0.5168). Insets show linear regression p-values.

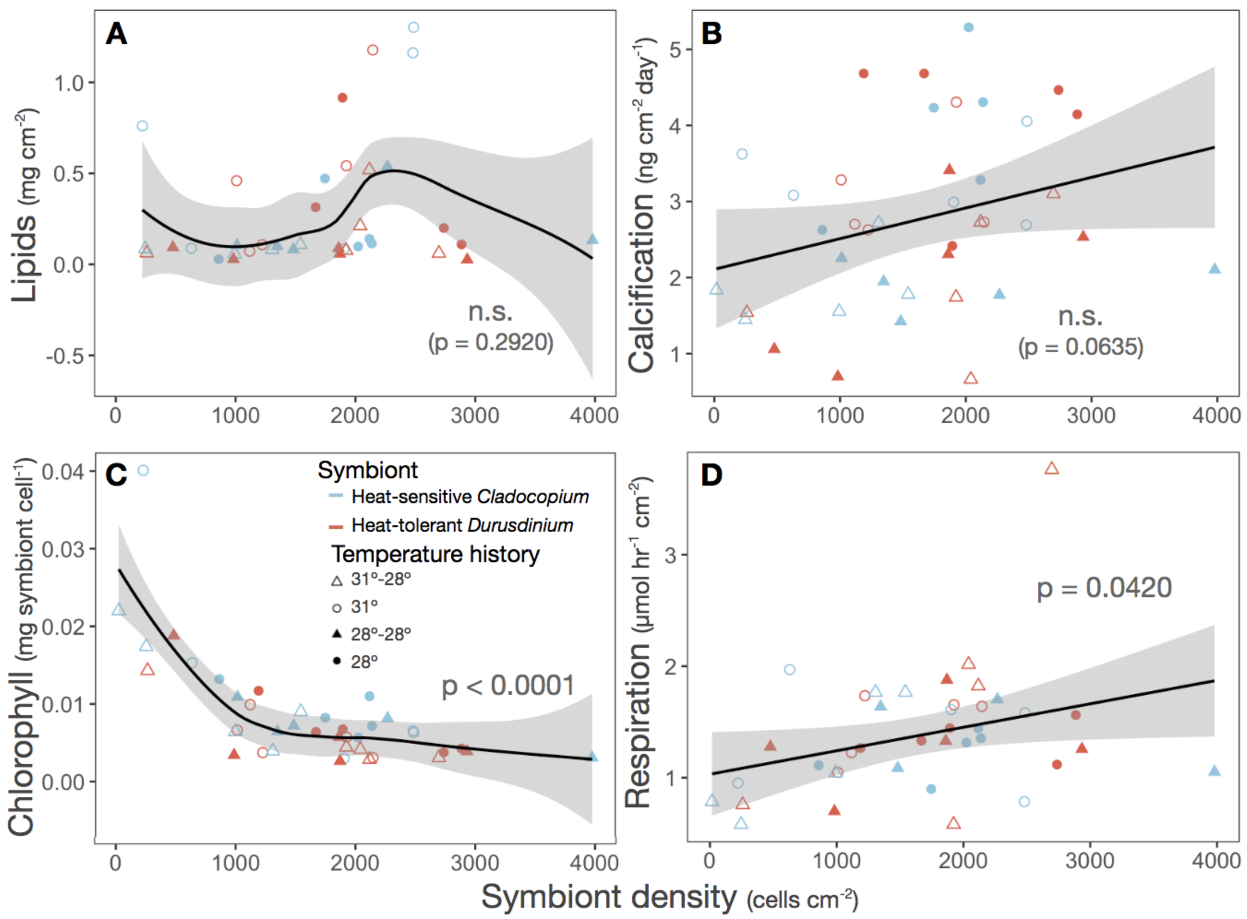


**Figure S8.** Relationship between symbiont density and host colony physiology. A) Symbiont density does not predict coral lipid content. Inset shows result of generalized additive model run using *gam* function in R package *mgcv*, method = REML (edf = 2.571, F = 1.193, REML score = 16.5851, p = 0.2920). B) Symbiont density does not predict coral calcification rate. Inset shows result of linear regression (t = 1.9113, df = 38, $R^2$ = 0.0877, p = 0.0635). C) Chlorophyll per symbiont decreases with symbiont density. Inset shows result of generalized additive model run using *gam* function in R package *mgcv*, method = REML (edf = 1.06, F = 1.001e+32, REML score = -777.856, p < 0.0001). D) Respiration increases with symbiont density. Inset shows result of linear regression (t = 2.1064, df = 37, $R^2$ = 0.1071, p = 0.04201).

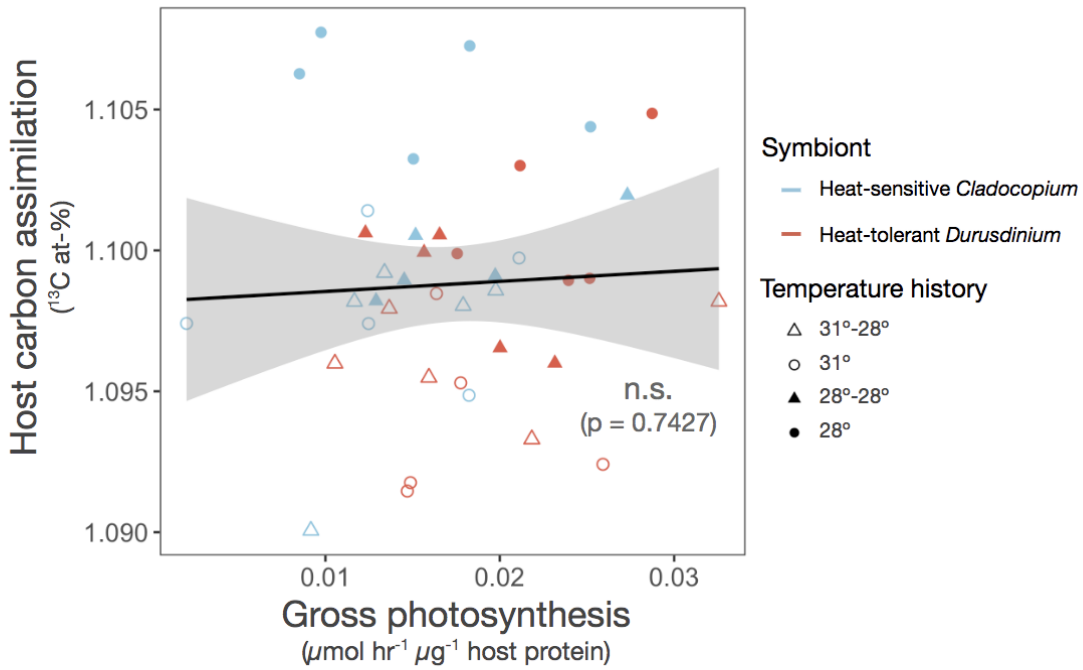


**Figure S9.** Gross photosynthesis does not predict host photosynthate assimilation ($^{13}C$ atom-%). Inset shows result of linear regression ($t = 0.33062$, $df = 38$, $R^2 = 0.00287$, $p = 0.7427$).

**Table S2** (included as separate Excel file). Results of statistical analyses testing effect of temperature on symbiont and coral colony physiological performance subsetted by experimental timepoint using linear mixed-effects models. Random intercept = genet. Model class = lmer. Significant effects determined by Satterthwaite's Type III ANOVA ($p<0.05$) indicated in bold. Abbreviations: df, degrees of freedom (Num, Den).

**Table S3** (included as separate Excel file). Results of statistical analyses testing effect of experimental timepoint on symbiont and coral colony physiological performance subsetted by temperature using linear mixed-effects models. Random intercept = genet. Model class = lmer. Significant effects determined by Satterthwaite's Type III ANOVA ($p<0.05$) indicated in bold. Abbreviations: df, degrees of freedom (Num, Den). Where indicated, heavy-tailed data were Gaussianized using R package *LambertW* [1, 2].
